# Genome mining of coastal cenote sediment-associated *Streptomyces* sp. NCA360 strain uncovers novel biosynthetic gene clusters and their regulatory architecture

**DOI:** 10.64898/2026.09.08.750260

**Authors:** Perla A. Contreras-de la Rosa, Jorge H. Ramírez-Prado, Elsa Góngora-Castillo, Alejandra Prieto-Davó

## Abstract

The rise of antibiotic resistance has intensified the search for novel antimicrobial compounds, and bioprospecting of natural products from underexplored environments remains an effective strategy. The genus *Streptomyces* is one of the most prolific sources of pharmacologically active secondary metabolites, whose biosynthetic information is encoded in biosynthetic gene clusters (BGCs). However, many BGCs remain transcriptionally silent under standard laboratory conditions, underscoring the need to characterize the regulatory mechanisms governing their activation.

In this study, we sequenced and analyzed the genome of *Streptomyces* sp. NCA360. The strain was isolated from sediments of the coastal cenote (a natural sinkhole) in the Yucatán Peninsula and selected for its antimicrobial and enzymatic activities. The high-quality genome assembly (89.7% completeness, <1% contamination) encoded 26 BGCs, of which 11 showed low similarity to characterized clusters and were classified as putatively novel (nBGCs). Resistance-guided prioritization identified duplicated resistance determinants, including an additional glyceraldehyde-3-phosphate dehydrogenase copy within a PKS-II cluster and Biotin_lipoyl/Carboxyl_trans domains within two divergent NRPS/PKS-I clusters, that were classified as candidate chemotherapeutic gene clusters. Biosynthetic pathways associated with clinically relevant antibiotics, including monobactams, carbapenems, and cephalosporins, were also detected. The regulatory architecture of the nBGCs revealed 26 regulatory genes, 20 transcription factor binding sites, and 20 rare TTA codons, reflecting heterogeneous and often multifactorial regulatory schemes. *In silico* protein-protein interaction analysis further revealed a coordinated cross-cluster regulation. The analysis of *Streptomyces* sp. NCA360 genome expands our understanding of the biosynthetic and regulatory diversity of *Streptomyces* and highlights the potential of cenotes as unique environments and reservoirs of new bioactive compounds with pharmaceutical relevance.

## Introduction

Antibiotic resistance has emerged as one of the most pressing challenges to global public health, compromising the effectiveness of existing treatments and increasing the risk of infectious diseases worldwide. This crisis has intensified the need for novel therapeutic agents, including antimicrobials and anticancer compounds. Natural products remain among the most successful sources of clinically relevant drugs, and microbial secondary metabolites have historically contributed in the discovery of new antibiotics. Thus, the exploration of underexplored environments for microorganisms capable of producing novel bioactive compounds has become a major strategy in contemporary drug discovery [1–3]. Additionally, characterizing microorganisms from diverse environments can provide valuable insights into alternative mechanisms of bacterial survival during antibiotic exposure. Recent studies have suggests that antibiotic-susceptible populations can survive during antibiotic exposure without being genetically resistant by exchanging essential proteins through membrane vesicles [4]. Therefore, continuing the genomic exploration of microorganisms is essential to identify novel secondary metabolites with antimicrobial potential and to understand complex biological mechanisms that enable bacteria to withstand antibiotic stress.

Members of the phylum *Actinomycetota* are Gram-positive bacteria widely distributed across aquatic and terrestrial ecosystems. These microorganisms play important ecological roles in organic matter decomposition and are recognized for their remarkable capacity to synthesize structurally diverse secondary metabolites [5–7]. Among them, the genus *Streptomyces* stands out for its ability to produce nearly two-thirds of all known antibiotics, making it key to drug discovery and development [5,8]. Predominantly associated with soil environments, *Streptomyces* species represent one of the most extensively studied bacterial groups for the identification of bioactive molecules, including compounds with antifungal, antiparasitic, immunosuppressive, and anticancer activities [9–11].

The extraordinary biosynthetic potential of *Streptomyces* is largely associated with their large genomes, which typically range from 4.5 to more than 11.5 Mb and encode numerous biosynthetic gene clusters (BGCs) responsible for specialized metabolite biosynthesis [12,13]. Recent comparative genomic studies have shown that the *Streptomyces* pangenome is open, indicating that its genetic repertoire continues to expand as additional genomes are sequenced and analyzed. This evolutionary pattern is thought to reflect frequent gene gain, gene loss, and horizontal gene transfer events, which contribute to the diversification of specialized metabolic pathways across the genus [14]. Thus, strains sharing nearly identical taxonomic marker genes, such as 16S rRNA or rpoB, may differ considerably in both the number and composition of their BGCs. Therefore, genome sequencing and analysis of individual strains provide valuable insights into strain-specific biosynthetic potential that may not be predicted solely from species- level classifications or previously sequenced representatives [15].

The increasing availability of microbial genome sequences has transformed natural product discovery by enabling genome mining approaches for the identification of BGCs with the potential to encode novel specialized metabolites [3,16–18]. Similarly, genome- guided analyses have facilitated the identification of compounds such as Abyssomicins from *Streptomyces* sp. LC-6-2 and Niphimycins from *Streptomyces* sp. IMB7-145, both of which exhibit promising antimicrobial activities [19,20].

A major challenge in natural product discovery is that many BGCs remain transcriptionally silent under standard laboratory conditions, limiting the exploitation of their biosynthetic potential [21]. Consequently, understanding the regulatory mechanisms controlling BGC expression has become an important aspect of genome mining and the functional characterization of specialized metabolism. BGC expression is typically controlled through hierarchical regulatory networks involving cluster-situated regulators and global regulatory systems that integrate environmental and developmental signals [22]. Many of these regulatory elements can be located within intergenic regions (IGRs). Although coding regions are often conserved among related strains, variation within IGRs may contribute to differences in regulatory patterns [14]. Furthermore, protein-protein interaction (PPI) analyses of BGC-associated regulatory proteins can provide insights into potential functional relationships among pathway-specific and global regulators [23–25]. Translational regulation also plays an important role in *Streptomyces*. The rare TTA leucine codon, whose translation depends on the bldA-encoded tRNA, has been associated with morphological differentiation and secondary metabolite production [26,27]. Therefore, the characterization of regulatory genes, TFBS, PPI networks, and rare TTA codons provides valuable insights into the regulatory potential of BGCs and their capacity for specialized metabolite biosynthesis.

Advances in genome mining have also enabled the prioritization of BGCs with potential pharmaceutical relevance. Recent efforts have focused on identifying chemotherapeutic gene clusters (CGCs), a group of BGCs predicted to encode compounds with antitumoral or other clinically relevant activities [15]. These approaches facilitate the prioritization of candidate clusters for downstream functional characterization and experimental validation, thereby supporting the discovery of natural products with potential medical applications.

In this context, the cenotes of the Yucatán Peninsula, Mexico, represent valuable and underexplored environments for microbial bioprospecting. These groundwater ecosystems are part of an extensive karst aquifer system characterized by unique physicochemical conditions, ecological isolation, and diverse microbial communities adapted to specialized niches [28,29]. Cenote sediments harbor complex microbial assemblages involved in biogeochemical processes and represent potential reservoirs of unexplored metabolic diversity [30,31].

In this study, we present the genome sequencing and genomic characterization of *Streptomyces* sp. NCA360, a strain isolated by Wissner *et al.* (2024) from sediments collected in the coastal cenote Pol-Ac (a Mayan term meaning “turtle head”) in the Yucatan Peninsula (S1 Fig). This strain was selected for genome sequencing and biosynthetic potential assessment based on its preliminary antimicrobial and enzymatic activities, as well as the PCR-based detection of type I polyketide synthase (PKS-I) genes, frequently associated with pharmacologically relevant molecules [31]. Genome mining revealed a diverse repertoire of biosynthetic gene clusters, including putative novel BGCs (nBGCs) with the potential to encode structurally distinct secondary metabolites. Furthermore, we explored how these clusters may be controlled and coordinated within the genome by characterizing their associated regulatory and translational features. Thus, we revealed that *Streptomyces* sp. NCA360 genome is a potential source of novel bioactive compounds. Additionally, this study emphasizes the value of genome-guided bioprospecting of microorganisms from unique ecosystems such as cenotes, and the importance of their conservation.

## Materials and Methods

### Bacterial strain

*Streptomyces* sp. NCA360 was previously isolated from sediment collected in the coastal cenote Pol-Ac, located within the El Palmar Ecological Reserve, Yucatán, Mexico. Details of sample collection, strain isolation, and its preliminary characterization have been reported by Wissner *et al.* (2024) [31] (S1 Fig). Based on its antimicrobial and enzymatic activities and the PCR-based detection of type I polyketide synthase (PKS-I) biosynthetic genes, the strain was selected for whole-genome sequencing and subsequent genomic analyses in this study.

### DNA isolation and genome sequencing

Genomic DNA was extracted using Zymo’s Quick-DNA Fungal/Bacterial Miniprep Kit (Cat. D6005) and subsequently purified using Promega’s Wizard® SV Gel and PCR Clean-Up System. DNA quantity and purity were assessed using a Nanodrop One spectrophotometer (Thermo Scientific, Waltham, USA) [32], considering a minimum concentration value of 150 ng/µl. DNA integrity was verified by electrophoresis in 1% agarose gel. A DNA library was constructed using Illumina MiSeq Reagent V3 following the kit instructions [33].

The genomic libraries were amplified by PCR and sequenced by shotgun. DNA sequencing was performed on the Illumina Miseq platform in paired-end (PE) mode, with a 150 bp read length and aiming for a ∼90x coverage, at the Centro de Investigación y de Estudios Avanzados del IPN (Cinvestav), Merida Unit, Yucatan, Mexico.

### Genome assembly and quality assessment

The quality of the genomic sequences was assessed using FastQC v.0.11.5 [34]. Low-quality reads (Phred < 20) and adapter sequences were removed with Trimmomatic v.0.39 [35] using the SLIDINGWINDOW function. Genome assembly was performed with the MaSuRCA v.4.1.0 program [36] in PE mode. Contigs were merged into supercontigs using Sequencher v.5.2.4 [37] with the “Dirty Data” (rigorous) assembly algorithm, applying a minimum overlap of 100 bp and a minimum identity of 95%. This procedure was performed independently three times, and any ambiguously joined supercontigs were subsequently separated.

The quality and integrity of the assembled genome were evaluated using the QUAST v.5.2.0 [38] and BUSCO v.5.7.1 [39] programs, using the bacteria_odb10 and streptomycetales_odb10 databases as references. Genome integrity and contamination were estimated using CheckM v.1.0.18 [40], employing the library of universal single-copy marker genes present in bacterial genomes. The GenoVi v.0.4.3 program was used to visualize the assembled genome [41].

### Taxonomic and functional genome annotation

The taxonomic classification of the assembled genome and digital DNA-DNA hybridization (dDDH) analysis were performed using the Type Strain Genome Server [42].

Phylogenetic analysis was performed using autoMLST: Automated Multi-Locus Species Tree (https://automlst.ziemertlab.com/), under the De novo option, employing IQ-TREE Ultrafast Bootstrap with 1000 replicates [43]. The phylogenetic tree was visualized and annotated using Interactive Tree of Life (iTOL; https://itol.embl.de/) [44].

Functional annotation of the genome was performed using Prokka v.1.14.6 with default parameters [45].

### Genome mining of biosynthetic gene clusters and metabolic pathway annotation

The identification of biosynthetic gene clusters (BGCs) associated with secondary metabolite production was performed using antiSMASH v.7.1.0 [46], in relaxed mode, allowing the detection of both complete and partial BGCs.

Putatively novel BGCs (nBGCs) were identified using BiG-SLiCE [47] in combination with the BiG-FAM v.1.0.0 database [48]. Clusters with a Euclidean distance >900 were classified as such according to the recommended threshold.

To prioritize BGCs with potential pharmaceutical relevance, the genome was analyzed using the Antibiotic Resistant Target Seeker (ARTS) 2.0 web server (https://arts.ziemertlab.com) [49] with default parameters. Metabolic pathway annotation was performed using BlastKOALA (http://www.kegg.jp/blastkoala/) [50], against the KEGG database [51] employing the genus_prokaryotes dataset for the functional annotation of predicted proteins.

### Analysis of regulatory elements and protein-protein interactions in nBGCs

For the analysis of regulatory elements within nBGCs, including regulatory genes, rare codons, and TFBS, clusters were selected based on their high dissimilar to known BGCs producing characterized secondary metabolites.

Regulatory genes were identified using antiSMASH [46], from the genome annotation report, with particular attention to genes explicitly annotated under this category.

Potential TFBS were subsequently detected using the TFBS-finder module of antiSMASH [52], which ranks predictions based on experimentally characterized binding sites in *Streptomyces* spp. genomes; only predictions with strong or medium confidence levels were retained for this analysis. In addition, the presence of the rare TTA leucine codon was identified within each nBGC using the same antiSMASH report.

Finally, PPI analysis was performed using FlashPPI [53], with the protein sequences encoded by each gene across the 11 nBGCs as input and a confidence threshold of 0.50. The analysis was subsequently focused on interactions involving those encoded by regulatory genes identified in the previous steps.

## Results

### Genome sequencing and assembly of *Streptomyces* sp. NCA360

Genomic sequencing of the *Streptomyces* sp. NCA360 generated a total of 2 million paired-end reads, after filtering 1.89 million (94%) were retained and assembled into 74 long supercontigs, with a genome size of 6.84 Mb, an N50 of 144,372 nt, and a maximum contig length of 505,256 nt (Table 1), and a GC content of 72.42% (Fig 1).

**Fig 1.**
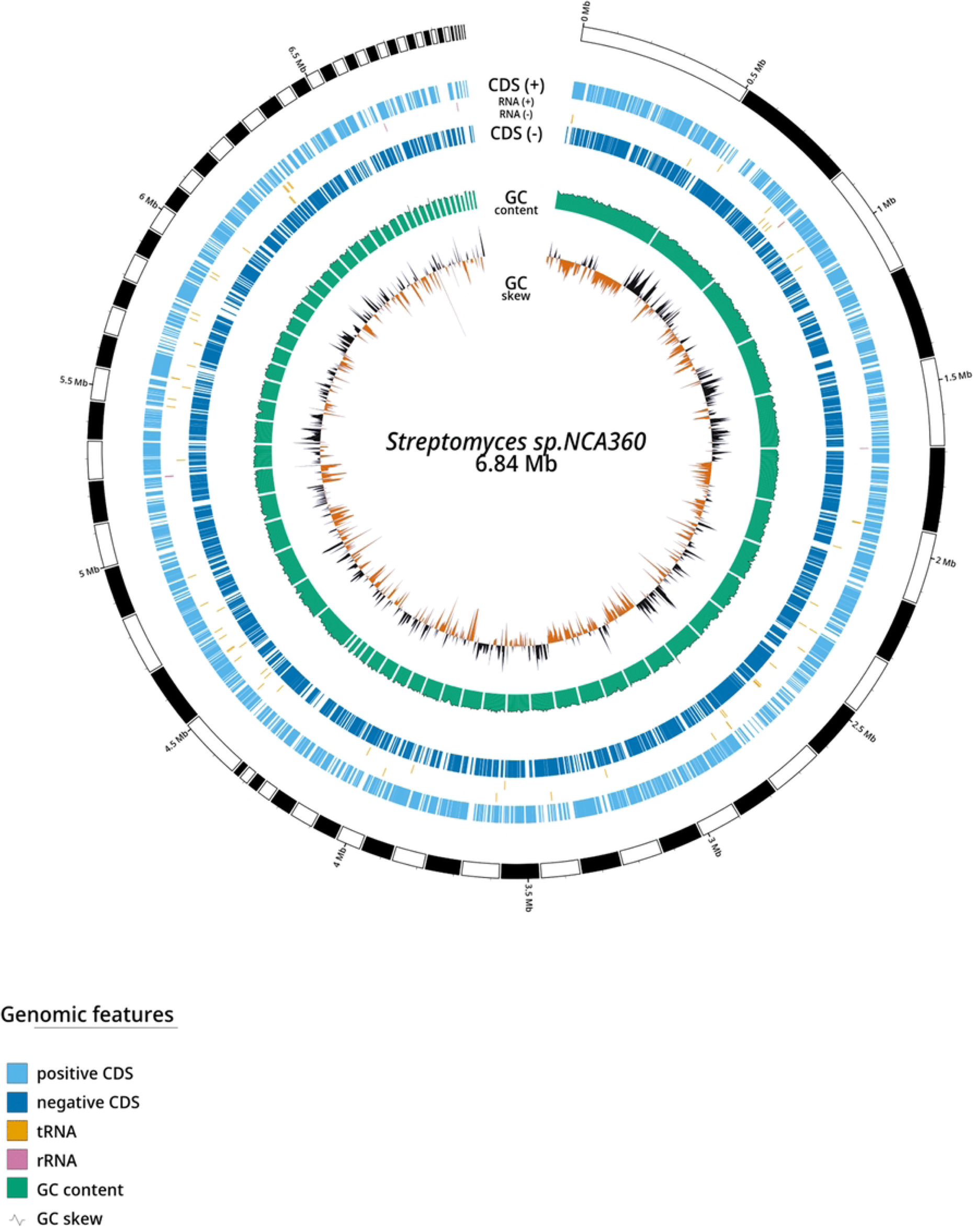
Circular visualization of the draft genome assembly of *Streptomyces* sp. NCA360. The assembly consists of 74 supercontigs displayed as alternating black and white segments, together with coding sequences (CDSs), tRNAs, rRNAs, and GC content. Supercontigs are arranged clockwise from the largest (505,256 bp) to the smallest (303 bp). Their arrangement is based solely on contig size and does not reflect their actual genomic order, orientation, or the unknown gaps between adjacent supercontigs.

**Table 1.**
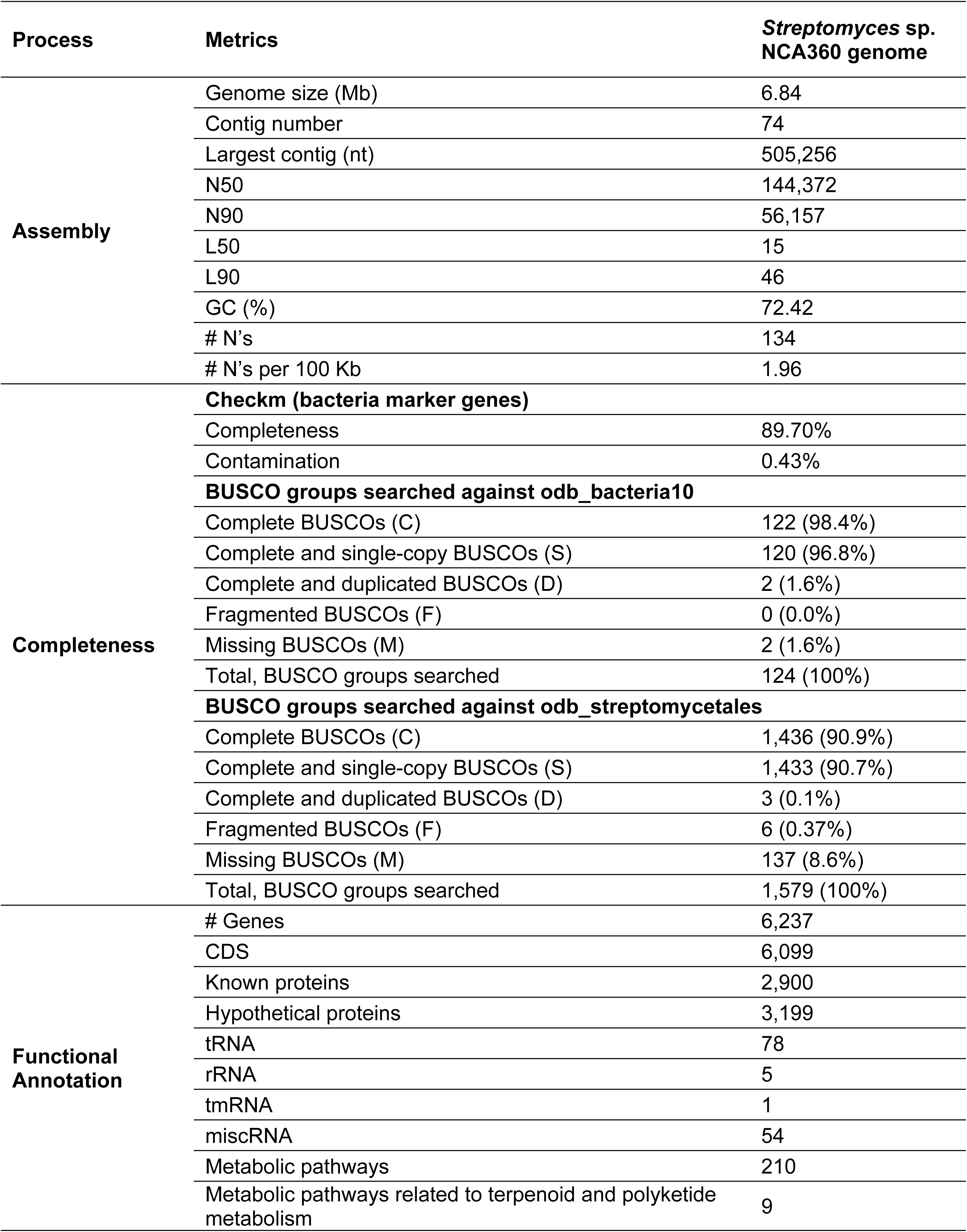
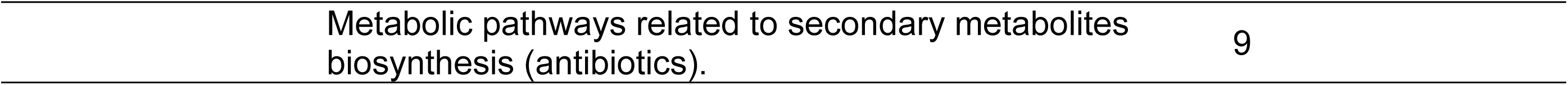
Genome assembly statistics and completeness assessment of *Streptomyces* sp. NCA360.

| Process | Metrics | *Streptomyces* sp.<br>NCA360 genome |
| --- | --- | --- |
| **Assembly** | Genome size (Mb) | 6.84 |
|  | Contig number | 74 |
|  | Largest contig (nt) | 505,256 |
|  | N50 | 144,372 |
|  | N90 | 56,157 |
|  | L50 | 15 |
|  | L90 | 46 |
|  | GC (%) | 72.42 |
|  | # N's | 134 |
|  | # N's per 100 Kb | 1.96 |
| **Completeness** | **Checkm (bacteria marker genes)** |  |
|  | Completeness | 89.70% |
|  | Contamination | 0.43% |
|  | **BUSCO groups searched against odb_bacteria10** |  |
|  | Complete BUSCOs (C) | 122 (98.4%) |
|  | Complete and single-copy BUSCOs (S) | 120 (96.8%) |
|  | Complete and duplicated BUSCOs (D) | 2 (1.6%) |
|  | Fragmented BUSCOs (F) | 0 (0.0%) |
|  | Missing BUSCOs (M) | 2 (1.6%) |
|  | Total, BUSCO groups searched | 124 (100%) |
|  | **BUSCO groups searched against odb_streptomycetales** |  |
|  | Complete BUSCOs (C) | 1,436 (90.9%) |
|  | Complete and single-copy BUSCOs (S) | 1,433 (90.7%) |
|  | Complete and duplicated BUSCOs (D) | 3 (0.1%) |
|  | Fragmented BUSCOs (F) | 6 (0.37%) |
|  | Missing BUSCOs (M) | 137 (8.6%) |
|  | Total, BUSCO groups searched | 1,579 (100%) |
| **Functional Annotation** | # Genes | 6,237 |
|  | CDS | 6,099 |
|  | Known proteins | 2,900 |
|  | Hypothetical proteins | 3,199 |
|  | tRNA | 78 |
|  | rRNA | 5 |
|  | tmRNA | 1 |
|  | miscRNA | 54 |
|  | Metabolic pathways | 210 |
|  | Metabolic pathways related to terpenoid and polyketide metabolism | 9 |

Quality assessment with CheckM [40], based on bacterial marker genes, indicated 89.7% completeness and less than 1% contamination. BUSCO analysis using bacterial orthologs (BUSCO-odb_bacteria10) and *Streptomycetales-*specific orthologs (BUSCO- odb_streptomycetales) [39], showed completeness levels of 98.4% and 90.9%, respectively (Table 1 and S2 Fig). According to the criteria of the Genome Standards Consortium (GSC) [54], these completeness values correspond to a high-quality genome assembly. The size and GC content of the assembled genome fall within the ranges reported for the genus *Streptomyces*, whose genomes typically range from 4.5 to12.6 Mb and have a GC content greater than 70% [13,17], consistent with the genomic characteristics commonly described for members of this genus.

Digital DNA-DNA hybridization (dDDH) analysis performed with the Type Strain Genome Server [42] revealed high genomic similarity between *Streptomyces* sp. NCA360 and several type species of the genus. dDDH values ranged from 77.8% to 85.0%, with differences in GC content of less than 0.2% (S1 Table). The highest similarity was observed with *S. variabilis* JCM 4422 (85.0%), *S. griseoincarnatus* JCM 4381 (84.1%), *S. erythrogriseus* JCM 9650 (83.2%), and *S. labedae* JCM 9381 (81.6%). All values exceed the commonly used genomic species boundary threshold (≥70%), suggesting intraspecific variation potentially associated with ecological adaptation.

Functional annotation revealed a total of 6,237 genes, including five rRNA genes (one 16S and four 5S), one tmRNA, 78 tRNA genes, 54 miscRNA genes, and 6,099 coding sequences (CDS), comprising 2,900 proteins with known functions and 3,199 hypothetical proteins (Fig 1, Table 1 and S2 Table).

Using the KEGG database [51], 210 metabolic pathways were annotated (Table 1), of which nine were related to terpenoid and polyketide biosynthesis (including ansamycins, enediynes, and tetracyclines), and ten corresponded to clinically relevant antibiotic biosynthesis pathways: (i) penicillins and cephalosporins, (ii) carbapenems, (iii) monobactams, (iv) streptomycin, (v) neomycin and kanamycin, (vi) gentamicin, (vii) carbose and validamycin, (viii) novobiocin, (ix) phenazine, and (x) prodigiosin (S3 Table). Additionally, coding genes of important industrial enzymes were identified, including three amylases, one cellulase, four chitinases, seven lipases, and 46 proteases (S2 Table), highlight the metabolic versatility and biotechnological potential of *Streptomyces* sp. NCA360.

### Biosynthetic potential of the NCA360 genome

Phylogenomic analysis placed *Streptomyces* sp. NCA360 in a well-supported clade (bootstrap value = 0.87) clustered with five type strains of the genus: *S. griseoincarnatus*, *S. althioticus, S. werraensis, S. cellulosae*, and *S. albogriseolus*, confirming its phylogenetic affinity with taxonomically validated representatives of the genus (Fig 2).

**Fig 2.**
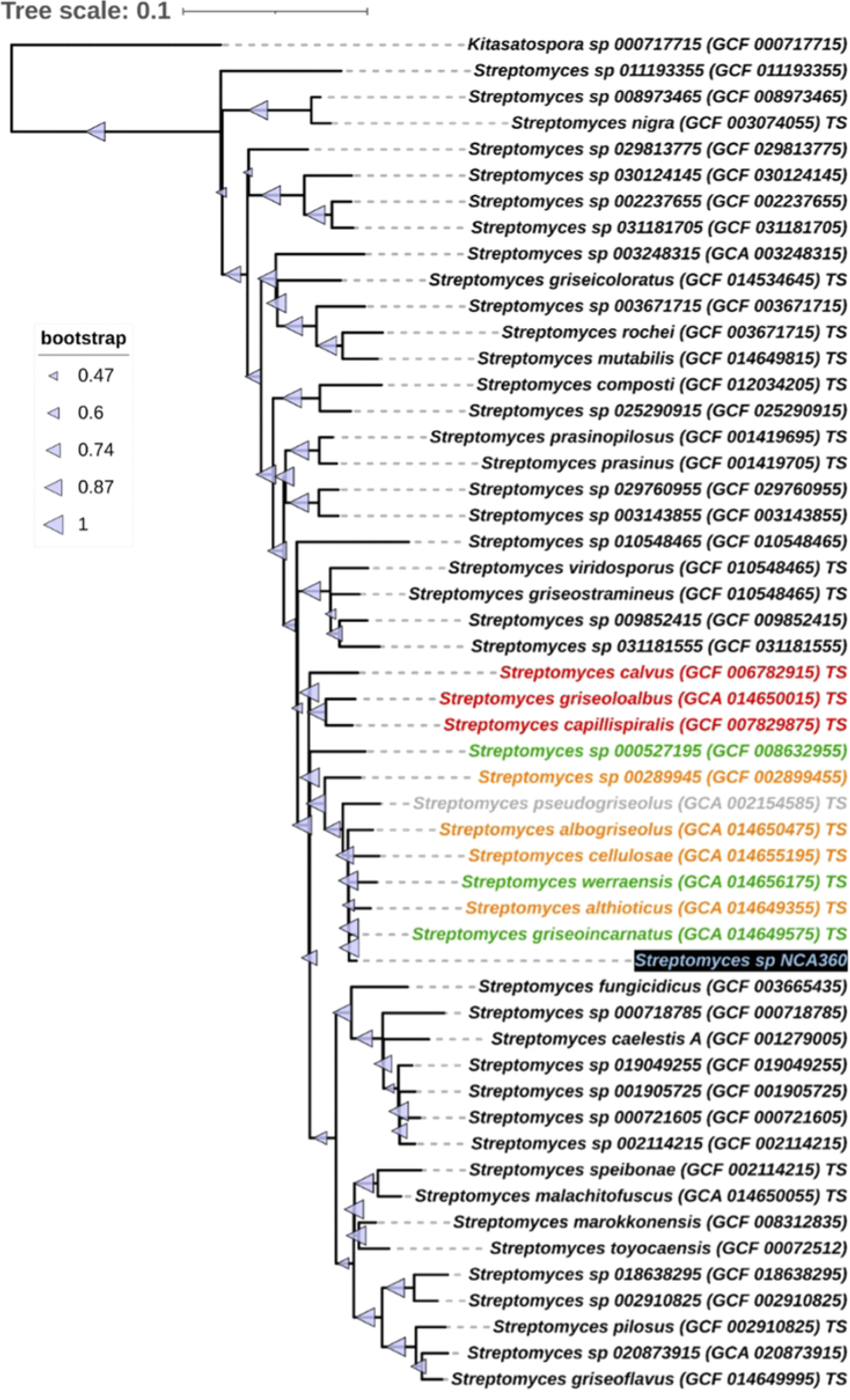
Multilocus phylogenetic tree showing the relationships between the genome of *Streptomyces* sp. NCA360 and representative *Streptomyces* strains. Bootstrap support values are indicated by triangles, with size proportional to the level of support. The position of strain NCA360 is highlighted in blue text on a black background. Label colors indicate the number of biosynthetic gene clusters (BGCs) identified per genome: green (<21 BGCs), orange (21–28 BGCs), red (>28 BGCs), and gray (no BGCs). Uncolored labels correspond to additional *Streptomyces* genomes included as phylogenetic references but not considered in the comparative BGC analysis. TS: genomes corresponding to type strains.

To explore biosynthetic potential of *Streptomyces* sp. NCA360, BGCs were identified and characterized by evaluating cluster diversity, genomic coverage and biosynthetic pathway types. A total of 26 BGCs were identified, spanning 797.76 Kb, and representing 11.6% of the assembled genome, with individual cluster sizes ranging from 4.7 Kb to 77.7 Kb.

The number of BGCs identified in the genomes of this clade ranged from 21 to 28, a range within which *Streptomyces* sp. NCA360 falls. Thus, the phylogenetic group as well as the BGC content suggest that the biosynthetic potential of NCA360 is comparable to that of closely related *Streptomyces* species.

The identified BGCs were classified into 16 different types: (i) betalactone, (ii) butyrolactone, (iii) ectoine, (iv) hydrogen-cyanide, (v) lanthipeptide, (vi) lassopeptide, (vii) non-ribosomal peptide-NRPS (NRPS), (viii) NRPS-Like, (ix) phenazine, (x) Polyketide synthase type I (PKS-I) and nonribosomal peptide synthetase (NRPS) hybrid (NRPS/PKS- I), (xi) Polyketide synthase type II (PKS-II), (xii) PKS-III, (xiii) ribosomal synthesized and post-translationally modified peptides (Ripp-Like), (xiv) RRE-containing, (xv) terpene and (xvi) NI-siderophore type. Of these, 14 were identified as complete clusters and 12 were located at contig edges. In terms of representation, NRPS/PKS-I hybrids were the most prevalent, with three clusters covering 2.99% of the genome and NRPS was the least abundant BGC type, with a single cluster accounting for 0.7% of the genome (S3 Fig, S4 Table).

### Identification of putative novel biosynthetic gene clusters

To identify novel BGCs (nBGCs), Euclidean distances were calculated and clusters with values greater than 900 were selected (37). A total of 11 putatively nBGCs were found, of which four were located within contigs and seven at contig edges (Table 2, S4 Fig). The 11 nBGCs were distributed across seven biosynthetic types: (i) ectoine, (ii) hydrogen-cyanide, and (iii) lanthipeptide, each represented by a single BGC, while (iv) NI-siderophore, (v) NRPS, (vi) NRPS/PKS-I, and (vii) PKS-II each contained two BGCs (Table 2).

**(i) Ectoine:** An ectoine-type nBGC was identified in Contig_7 (Euclidean distance= 994) consisting of 14 genes, of these two to four showed similarities of 33% to 100% with genes from previously described ectoine BGCs, while the remaining ten exhibited divergences from other known clusters, including butyrolactone, lactonamycin, and griseoviridin/fijimycin types.
**(ii) Hydrogen-cyanide:** A hydrogen-cyanide-type nBGC was identified in Contig_27 (Euclidean distance= 982) consisting of 12 genes, four of which share 28% similarity with a RiPP-type BGC associated with the metabolite aborycin.
**(iii) Lanthipeptide:** A lanthipeptide-type nBGC was identified in scf7180000000323 (Euclidean distance= 1,111), comprising 28 genes, four of which showed 100% identity with genes from two previously described BGCs encoding venezuelin, and 11% identity with an 18-gene BGC associated with valinomycin production.
**(iv) NI-siderophore:** A Ni-siderophore-type nBGC was detected in contig scf7180000000298 (Euclidean distance = 1,298), comprising 26 genes, six of which showed 100% identity to genes within a previously characterized BGC involved in desferrioxamine B/E biosynthesis. A second Ni-siderophore-type nBGC was identified in contig Contig_19, displaying 13% similarity to a previously characterized paulomycin-associated BGC. This cluster comprises 30 genes, six of which share 13% identity with genes from the reference cluster.
**(v) NRPS, (vi) NRPS/PKS-I, and (vii) PKS-II types:** Six nBGCs were identified, including two clusters of each biosynthetic type. NRPS-type nBGCs comprised 14 and 45 genes, with 11 genes showing similarity to previously characterized clusters, displaying 39% and 100% similarity and Euclidean distances of 1,219 and 934, respectively. NRPS/PKS-I hybrid nBGCs contained 50 and 37 genes, with 18 and 17 similar genes, respectively, showing 64% and 48% similarity and Euclidean distances ranging from 1,825 to 1,848. PKS-II nBGCs consisted of 59 and 77 genes, with 16 and 10 similar genes, respectively, exhibiting 88% and 83% similarity and Euclidean distances of 1,448 and 1,022. Overall, these clusters showed variable levels of similarity to previously characterized BGCs, suggesting differences in their biosynthetic architectures.

**Table 2.** Classification and characteristics of nBGCs in the genome of *Streptomyces* sp. NCA360.

| BGC type | Number of genes | Contig ID | Likely metabolite | Similarity (%) | Number of similar genes | Size (Kb) | Euclidean distance |
| --- | --- | --- | --- | --- | --- | --- | --- |
| Ectoine | 14 | Contig_7 | Ectoine | 100 | 4 | 14.3 | 994 |
| Hydrogen-cyanide | 12 | Contig_27 | Aborycin | 28 | 4 | 12.8 | 982 |
| Lanthipeptide | 28 | scf7180000000323 | Venezuelin | 100 | 4 | 36.2 | 1,111 |
| NI-siderophore | 26 | scf7180000000298 | Desferrioxamin B, Desferrioxamine E | 100 | 6 | 29.7 | 1,298 |
|  | 30 | Contig_19 | Paulomycin | 13 | 6 | 31.2 | 1,005 |
| NRPS | 14 | Contig_30 | Naphthyridinomycin | 39 | 11 | 16.0 | 1,219 |
|  | 45 | Contig_3 | Coelibactin | 100 | 11 | 63.4 | 934 |
| NRPS/ PKS-I | 50 | Contig_3 | Naphthyridinomycin | 64 | 18 | 77.7 | 1,848 |
|  | 37 | scf7180000000260 | Polyoxypeptin | 48 | 17 | 74.2 | 1,825 |
| PKS-II | 59 | Contig_24 | Resistomycin, resistoflavine | 88 | 16 | 69.9 | 1,448 |
|  | 77 | Contig_20 | Spore pigment | 83 | 10 | 72.5 | 1,022 |

These 11 nBGCs exhibited diverse patterns of similarity to known biosynthetic clusters, ranging from conserved regions to highly divergent architectures and underscores the potential of these nBGCs to encode structurally distinct secondary metabolites.

### Resistance-guided prioritization of novel biosynthetic gene clusters and horizontal gene transfer analyses

To prioritize nBGCs with potential pharmaceutical relevance [15], we analyzed the genome using ARTS 2.0 [49], a genome-mining framework that identifies duplicated essential genes, horizontal gene transfer (HGT) events, and resistance models (ResModels) associated with secondary metabolite biosynthesis. A total of 36 ResModel hits and 57 duplicated core genes were identified, indicating the presence of multiple potential self-resistance mechanisms associated with secondary metabolism (S5 and S6 Table). In addition, 16 genes showing evidence of horizontal transfer were identified across 18 of the 26 BGCs present in the genome (S7 Table). This distribution suggests that horizontal gene acquisition may have contributed to the diversification of the strain’s biosynthetic repertoire [49].

Out of the 11 nBGCs, three contained ResModels (Fig 3). The type II polyketide synthase (PKS-II) nBGC located on Contig_20 contained a copy of the gene encoding glyceraldehyde-3-phosphate dehydrogenase (GAPDH), which was predicted as a potential resistance determinant (Fig 3). This nBGC was annotated as a biosynthetic pathway associated with spore pigments. The other two nBGCs containing ResModels corresponded to highly divergent hybrid NRPS/PKS-I clusters. The nBGC associated with naphthyridinomycin, located on Contig_3, contained a gene encoding a Biotin_lipoyl domain, whereas the polyoxypeptin-related nBGC, located on scf7180000000260, contained a gene encoding a Carboxyl_trans domain (Fig 3).

**Fig 3.**
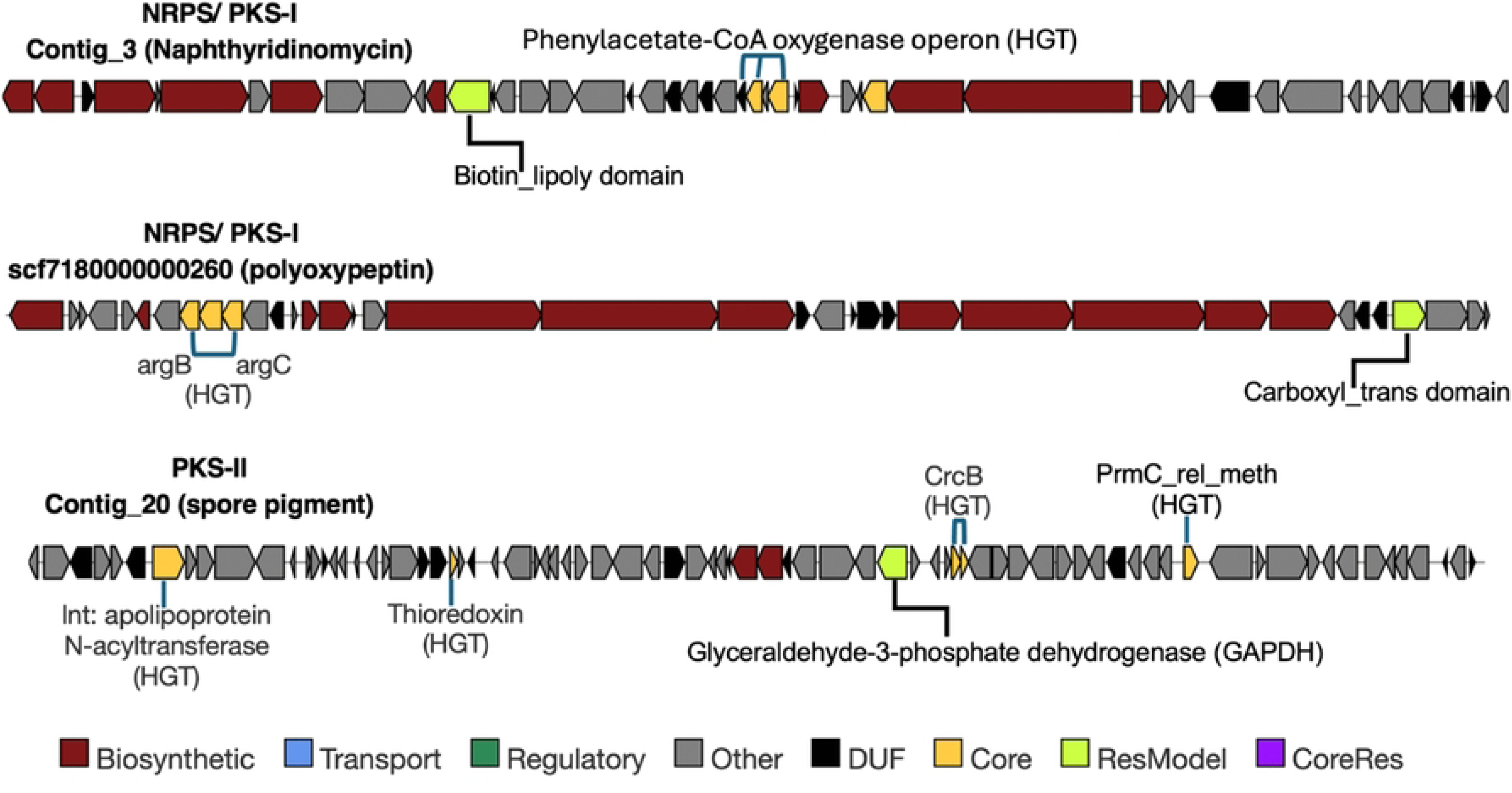
ARTS-prioritized novel biosynthetic gene clusters (nBGCs). Composite representation of the three nBGCs identified by ARTS 2.0 as containing resistance models. The figure highlights resistance-associated features, as well as genes showing evidence of horizontal gene transfer (HGT). Functional categories are indicated according to the color legend.

Evidence of HGT was particularly relevant among these prioritized nBGCs. Seven of the 11 nBGCs contained genes showing evidence of HGT, including those three nBGCs with ResModels (Fig 3; S8 Table). Among these, the naphthyridinomycin-associated hybrid NRPS/PKS-I nBGC located on Contig_3 had the highest number of genes showing evidence of HGT. These included an operon comprising three genes encoding components of the phenylacetate-CoA oxygenase pathway (PA_CoA_Oxy1, PA_CoA_Oxy3, and PA_CoA_Oxy4) (Table S5; Fig 3). The nBGC located on Contig_20 contained the HGT-associated genes of thioredoxin, lnt, PrmC_rel_meth, and two copies of crcB, whereas the polyoxypeptin-related nBGC on scf7180000000260 contained argB and argC with evidence of HGT (S8 Table).

Additionally, copies of several essential genes associated with translation showed evidence of HGT, including the gene encoding ribosomal protein S18, rpmF, and elongation factor G (EF-G), which were located in proximity to biosynthetic regions (S7 Table). These findings suggest that HGT is not restricted to genes directly associated with biosynthesis or resistance but may also involve genes related to essential cellular processes, potentially contributing to the adaptation and evolution of the biosynthetic repertoire.

### Regulatory architecture of novel biosynthetic gene clusters and protein interaction networks underlying secondary metabolism in NCA360

Due to secondary metabolism being tightly regulated and frequently influenced by environmental conditions, we next explored the regulatory architecture of the putative nBGCs through the analysis of their regulatory features. Previous studies have shown that regulatory components associated with biosynthetic clusters can evolve rapidly in response to ecological specialization [14]. A targeted analysis of three regulatory-associated features was performed across the 11 nBGCs identified in the *Streptomyces* sp. NCA360 genome: (i) regulatory genes encoded within the clusters, (ii) transcription factor binding sites (TFBS), and (iii) TTA codons (Table 3 and S4 Fig). These regulatory features were identified in five out of seven nBGC classes detected in the genome.

**Table 3.** Information on regulatory elements identified in 11 nBGCs.

| BGC type | Novel BGCs | TTA codons | TFBS | Regulatory genes | Regulator gene type in BGC |
| --- | --- | --- | --- | --- | --- |
| Ectoine | 1 | 2 | NA | 3 | TetR family, SARP family (transcriptional regulators) |
| Hydrogen-cyanide | 1 | 2 | NA | 2 | GntR family, LacI family (transcriptional regulators) |
| Lanthipeptide | 1 | 4 | 1 | 1 | SARP family (transcriptional regulator) |
| **NI-siderophore** | 2 | 3 | 6 | 8 | TetR, AraC, LacI, IclR families (transcriptional regulators); Sensor histidine kinase (repressor); LuxR DNA-binding response regulator |
| **NRPS** | 2 | 4 | 6 | 3 | ArsR, MerR families (transcriptional regulators); Hydrogen peroxide sensitive (repressor) |
| **NRPS/ PKS-I** | 2 | 2 | 2 | 3 | SARP, MerR families (transcriptional regulators); RNA polymerase sigma-24 subunit (ECF subfamily) |
| **PKS-II** | 2 | 3 | 5 | 6 | TetR, MarR, GntR, PadR families (transcriptional regulators); LuxR DNA-binding response regulator |

A total of 26 regulatory genes were identified (Table 3 and S4 Fig). The NI-siderophore and PKS-II nBGCs contained eight and six genes, respectively, distributed across two clusters each. In NI-siderophore clusters, the identified genes corresponded to four different transcription factor families, in addition to a repressor and a binding protein, suggesting a multifactorial and potentially coordinated regulation. In PKS-II clusters, regulators related to four transcriptional families and one binding family were detected. In contrast, the Lanthipeptide-type nBGC contained a single regulator from the SARP family, a recognized transcriptional activator, while the remaining clusters showed a range of two to three regulators, reflecting varying levels of regulatory complexity.

A total of 20 TFBS were identified, approximately 15 bp in length, distributed across 11 categories (Table 3). The nBGCs with the highest number of TFBS were those of the NI- siderophore and NRPS types, with six each, followed by PKS-II with five. In NI-siderophore clusters, TFBS sequences corresponded to the DmdR1, CelR, BldD, AfsR, and OsdR categories, associated with iron and cellobiose uptake, global developmental regulation, antibiotic production, pleiotropic regulation, and stress response. Out of two NRPS identified clusters, only one contained six TFBS, all belonging to the ZuR family (zinc- sensitive repressor). Among the PKS-II clusters, five TFBS representing four different categories were identified, while the remaining nBGC types contained between zero and two TFBS (Table 3 and S4 Fig).

In addition, 20 TTA codons were identified, distributed across all seven nBGCs types, at a frequency of two to four per cluster (Table 3 and S4 Fig). These codons were found in genes with diverse functions including regulatory, transport, core biosynthetic, and accessory functions, suggesting that they constitute an additional layer of translational control capable of modulating expression efficiency across multiple functional categories. For instance, the Lanthipeptide and NRPS clusters harbored the highest number of TTA codons with four each. These results show that the nBGCs of *Streptomyces* sp. NCA360 exhibit heterogeneous regulatory architectures. Clusters such as NI-siderophore and PKS-II, stand out for the abundance and diversity of both regulatory genes and TFBS, while others, such as the lanthipeptide cluster, display comparatively simpler schemes that may be complemented by the presence of TTA codons.

Additionally, to investigate the functional connectivity among proteins encoded within the nBGCs, we performed protein-protein interaction (PPI) computational analyses. This approach enabled the identification of potential cross-regulatory interactions among distinct classes of BGCs and provided insights into how regulatory proteins may coordinate expression and activity of specialized metabolic pathways. The PPI analysis of the 393 protein sequences encoded across the 11 nBGCs (S10 Table) revealed that 73 are connected through 165 interactions and organized into 11 functional modules (Fig 4, and S11 Table).

**Fig 4.**
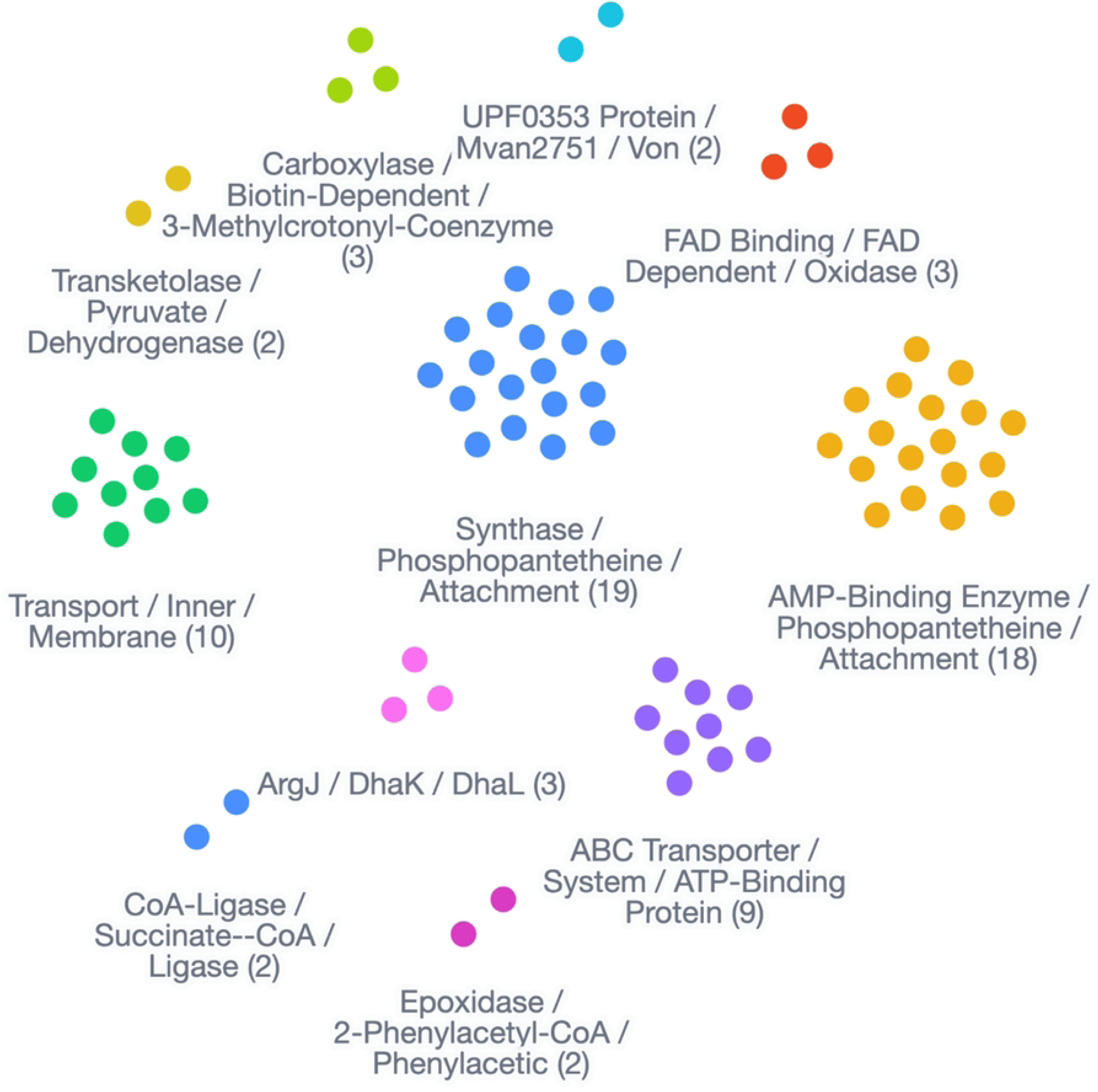
PPI network generated by FlashPPI for the proteins encoded within the nBGCs of *Streptomyces* sp. NCA360. The predicted network comprises 73 proteins connected through 165 interactions, organized into 11 functional modules. Each node represents a protein; the number of proteins per module is indicated in parentheses. A minimum contact score of 0.50 was applied.

To identify interactions involving regulatory proteins, the 26 regulatory genes previously identified were mapped onto the network, each represented by its corresponding protein sequence. Of these, four proteins were identified within the regulatory network. The sensor histidine kinase (nisiderophore_c19_ctg12_105) and the acyl carrier protein (pksii_c24_ctg23_29) were both connected to the isonitrile lipopeptide synthase (nrpspksi_c3_ctg2_4), which also interacted with the putative RNA polymerase sigma factor (nrpspksi_ctg2_37) (Fig 5A). These interactions revealed regulatory-associated proteins from different nBGC types, including NI-siderophore, NRPS/PKS-I, and PKS-II clusters, suggesting possible coordinated regulation among biosynthetic clusters.

**Fig 5.**
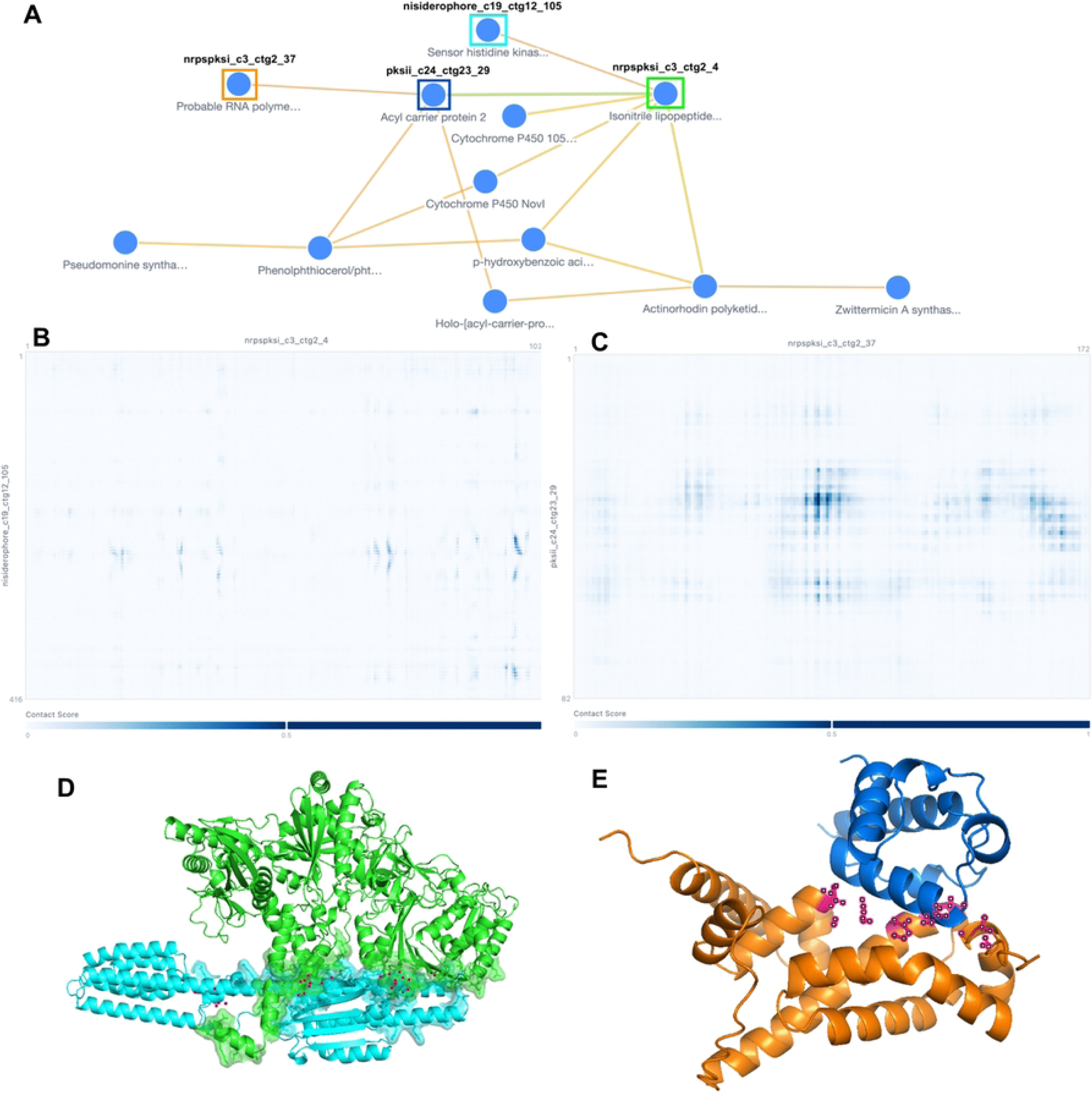
Protein-protein interaction computational analysis among regulatory genes identified in the nBGCs of *Streptomyces* sp. NCA360. (A) Interactions among four regulatory proteins from three nBGC types: NI-siderophore, NRPS/PKS-I, and PKS-II. (B, C) Residue-residue contact maps representing the probability of spatial proximity between residue pairs for the sensor histidine kinase/isonitrile lipopeptide synthase (B) and RNA polymerase sigma factor/acyl carrier protein (C) complexes. (D, E) Structural confidence metrics for each complex: chain_ptm = 0.65 and 0.38, ipTM = 0.17 (D); chain_ptm = 0.54 and 0.74, ipTM = 0.13 (E).

To further evaluate the potential interaction interfaces among these proteins, residue– residue contact maps were generated. These maps represent the predicted spatial proximity between pairs of residues within potentially interacting proteins (Fig 5B and C). Structural analyses were performed using representative protein models, including the sensor histidine kinase/isonitrile lipopeptide synthase complex (Fig 5B) and the predicted RNA polymerase sigma factor/acyl carrier protein interaction (Fig 5C). For the sensor histidine kinase/isonitrile lipopeptide synthase complex, individual protein models showed chain pTM values of 0.65 and 0.38, respectivel, whereas the predicted interface pTM (ipTM) value was 0.17 (Fig 5D). For the predicted RNA polymerase sigma factor/acyl carrier protein interaction, chain pTM values were 0.54 and 0.74, respectively, with an ipTM value of 0.13 (Fig 5E). In both cases, the individual protein models displayed moderate folding confidence, while the low ipTM values indicate limited confidence in the predicted protein–protein interfaces, suggesting that these interactions require further experimental validation.

## Discussion

The continued emergence of multidrug-resistant pathogens has intensified the search for novel bioactive compounds. Members of the genus *Streptomyces* remain one of the most prolific sources of natural products. Although several of *Streptomyces* genomes have been sequenced during the last decade, comparative genomic studies have consistently demonstrated that the genus possesses an open pangenome, in which each newly sequenced strain contributes additional accessory genes, particularly BGCs [14]. The NCA360 strain was cultivated and selected for sequencing based on its hydrolytic activities, including amylase, cellulase, gelatinase, chitinase, and lipase [31]. After sequencing, our genomic analysis confirmed these traits by identifying the corresponding genes (S2 Table), reinforcing the strain’s biotechnological potential.

The genomic characterization of *Streptomyces* sp. NCA360 expands our understanding of the biosynthetic diversity harbored by microorganisms inhabiting the cenotes of Yucatán Peninsula, unique karst ecosystems. Genome size, GC content, and BGC repertoire were consistent with the ranges typically reported for the genus, which vary between 4.5 and 12.6 Mb, GC contents exceeding 70%, and 8–83 BGCs per genome [13,15,17]. The high-quality genome assembly enabled a comprehensive characterization of the strain’s biosynthetic repertoire. Large biosynthetic gene clusters, particularly hybrid NRPS/PKS systems, frequently contain repetitive domains and genes spanning several kilobases, making them difficult to assemble accurately [55]. Therefore, the high contiguity of the NCA360 assembly, together with its BUSCO completeness score of 98.4%, provides confidence that the identified biosynthetic gene clusters represent their genomic organization (Table 1 and S2 Fig).

Previous studies have demonstrated that strains sharing nearly identical marker gene sequences, including 16S rRNA and rpoB, can exhibit substantial differences in their biosynthetic repertoires [14]. Comparative genomic analyses revealed that although *Streptomyces* sp. NCA360 shares high genomic similarity with closely related *Streptomyces* species, it also possesses unique genomic features that suggest it may represent a specialized lineage adapted to the cenote environment. Furthermore, the number of predicted coding sequences (CDSs) is consistent with the genomic complexity reported for other *Streptomyces* genomes [55]. Many of these genes are likely to belong to the accessory genome, including the strain-specific fraction commonly referred to as the cloud genome, which has been associated with environmental adaptation and the diversification of specialized metabolism (13).

Previous PCR-based screening had suggested the presence of PKS-I biosynthetic genes in NCA360 [31]. However, whole-genome sequencing considerably expanded this observation by revealing a broader and more structurally diverse biosynthetic repertoire. The identification of 11 putatively nBGCs displaying high structural divergence, including ectoine, hydrogen-cyanide, NRPS/PKS-I hybrids, PKS-II, Lanthipeptide, and NI- siderophore types, strongly supports this model and suggests that these clusters constitute part of the cloud genome that enables adaptation to the unique physicochemical conditions of Pol-Ac cenote sediments [56]. Among the identified biosynthetic pathways, the hybrid NRPS/PKS-I clusters represent the most promising candidates for future functional characterization. Comparative analyses demonstrated that the two most divergent clusters share only limited similarity with known biosynthetic systems, despite exhibiting partial homology to polyoxypeptin- and Naphthyridinomycin- associated pathways (Table 2). Such structural divergence suggests that these clusters may encode previously undescribed biosynthetic pathways rather than simple variants of known metabolites. Hybrid NRPS/PKS systems are recognized as one of the richest sources of chemically complex natural products, frequently producing compounds with antimicrobial, immunosuppressive, or antitumor activities [15,57–59].

The NCA360 genome showed several duplicated ribosomal proteins. Duplicated essential genes and horizontal gene transfer events are two genomic signatures commonly linked to antibiotic production and self-resistance. Resistance genes are frequently mobilized together with biosynthetic gene clusters, allowing microorganisms to acquire both metabolite production and self-protection simultaneously [49]. In the NCA360 genome, 36 ResModels and 57 duplicated core genes, including S18, rpmF, and EF-G, indicate that several biosynthetic pathways are accompanied by intrinsic protection mechanisms that enable the bacterium to tolerate its own secondary metabolites [49,60].

Specifically, the novel PKS-II cluster located on Contig_20, which contains an additional copy of glyceraldehyde-3-phosphate dehydrogenase (GAPDH), suggests that this enzyme may function as a self-resistance determinant. Natural products capable of inhibiting GAPDH have previously been described [61–63] supporting the hypothesis that the additional GAPDH copy identified within this cluster may protect the producer organism while preserving central carbon metabolism during specialized metabolite biosynthesis. Although this evidence does not demonstrate that GAPDH is the molecular target of the metabolite encoded by this PKS-II cluster, it is consistent with self-resistance strategies frequently associated with specialized metabolite biosynthetic pathways. Beyond self-resistance, the additional GAPDH copy may also contribute to maintaining the glycolytic flux required to sustain polyketide biosynthesis by ensuring the continuous supply of central metabolic intermediates that generate precursors such as malonyl-CoA, an essential building block for PKS-II assembly. Additionally, this cluster was predicted to encode a spore pigment-like metabolite (Table 2), and several studies suggest that bacterial pigments are multifunctional secondary metabolites contributing to microbial competition through antimicrobial activity, protection against oxidative stress, ultraviolet radiation, and other environmental challenges [64].

Similarly, the resistance signatures detected within the putative naphthyridinomycin- and polyoxypeptin-associated clusters further increase their pharmaceutical relevance. These clusters contain resistance models involving the Biotin_lipoyl and Carboxyl_trans domains, which are associated with fatty acid metabolism and malonyl-CoA utilization [65,66]. Because both pathways have previously been associated with cytotoxic compounds, these nBGCs can be considered candidate chemotherapeutic gene clusters (CGCs), a recently proposed term describing biosynthetic clusters with potential anticancer applications [15]. The pharmaceutical potential of NCA360 is further supported by the presence of biosynthetic pathways with potential antimicrobial activity, particularly in the context of the urgent need for new agents against WHO-priority pathogens, including Acinetobacter baumannii and Klebsiella pneumoniae [67].

The presence of a complete phenylacetate degradation operon within the putative naphthyridinomycin/coelibactin genomic region (Contig_3) (Fig 3 and S8 Table) may represent an adaptive hotspot that enables NCA360 to exploit locally available substrates. The aromatic compounds derived from plant material constitute an important carbon source in organic-rich sediments. The acquisition of this operon through horizontal gene transfer suggests that some BGCs are not isolated biosynthetic units but rather components of larger adaptive genomic islands integrating metabolism, ecological specialization, and chemical defense [68].

Recent comparative genomic studies have shown that IGRs are rarely conserved across the *Streptomyces* genus and are generally maintained only among closely related species, highlighting the rapid evolutionary diversification of transcriptional regulation [14]. This was evident in our regulatory analyses in which multiple regulatory families, including TetR, GntR, LuxR, LacI, MerR, ArsR, SARP, and ECF sigma-associated regulators, as well as multiple predicted TFBSs, indicates that the nBGCs of *Streptomyces* sp. NCA360 are controlled by specialized regulatory networks (Table 3, S9 Table). Many of these regulators are associated with responses to nutrient availability, oxidative stress, metal homeostasis, quorum sensing, and developmental transitions, suggesting that secondary metabolite production is tightly coordinated with environmental sensing and cellular physiology [22]. The detection of TFBSs involved in metal homeostasis (Zur and Nur), DNA damage and oxidative stress responses (LexA, HypR, and OsdR) [69–76] and primary metabolic pathways (ArgR, CelR, NrdR, and NrtR) [77,78,73,72,79], supports the hypothesis that NCA360 biosynthetic pathways are embedded within broader adaptive networks that respond to the fluctuating physicochemical conditions characteristic of cenote sediments. An additional layer of regulation may be provided by rare TTA codons, which require the bldA-encoded tRNA for translation, and which have been linked to antibiotic biosynthesis and morphological differentiation [26,27,80,81]. This regulatory integration is further supported by the protein-protein interaction *in silico* analyses, which revealed interconnected networks among proteins encoded by different BGCs associated to siderophore, PKS-II, and hybrid NRPS/PKS-I pathways. The interaction between a sigma factor and an acyl carrier protein (ACP) establishes a direct connection between transcriptional regulation and metabolite assembly, suggesting that secondary metabolite production in NCA360 is not organized as a set of isolated pathways but is integrated into a broader metabolic and adaptative response [82,83].

While horizontal gene transfer is well recognized as an important mechanism for the spreads of antibiotic resistance genes, the mechanisms and effects of horizontal protein transfer remain considerably less understood. Recent work by Wen *et al.* (2026) [84] described a bacterial survival strategy in which antibiotic-stressed cells exchange essential proteins with neighboring bacteria through membrane vesicles, thereby promoting survival without requiring the acquisition of conventional antibiotic resistance genes. In this context, it is noteworthy that the genome of *Streptomyces* sp. NCA360 contains *pspA* (phage shock protein A), a essential gene for membrane vesicle production, as well as *relA*, a key gen for the stringent response that promotes survival under stress by reducing cellular metabolic activity (S9 Table). Although the presence of *pspA* and *relA* alone does not demonstrate the existence of a vesicle-mediated protein exchange system in NCA360, their association with stress adaptation and cellular persistence through horizontal protein transfer raises the possibility that this strain may employ survival strategies analogous to those described by Wen *et al.* (2026) [84], and represents an interesting direction for future research.

Finally, our work highlights the importance of bioprospecting in unique ecosystems such as cenotes, whose conservation is essential to preserve their microbial biodiversity and the vast chemical repertoire they offer for applications in health, industry, and science.

## Acknowledgments

The authors would like to express their gratitude to the technical staff at the Centro de Investigación Científica de Yucatán (CICY), Luis Alberto Puc Canul and Mirbella Cáceres, and the technical staff at the Facultad de Química, Universidad Nacional Autónoma de México (UNAM), Norma Angélica Márquez Velázquez and Dr. Wendy Itzel Escobedo Hinojosa. We also thank Illumina technical assistance staff Alejandra García, Valeria Ortega, Verónica Sánchez, Ileana Gutiérrez, Sandra Pérez, Abril Gamboa and Dr. José Q. García Maldonado from Centro de Investigación y de Estudios Avanzados del Instituto Politécnico Nacional (Cinvestav), Mérida. Special thanks to “Cenoteando” (https://cenoteando.mx/expediciones/), divers, technicians, and specialists, whose expertise and assistance were essential to the success of this project.

## Supporting information

**S1 Fig. Colony morphology of *Streptomyces* sp. NCA360.** Photograph showing the colony morphology of *Streptomyces* sp. NCA360 grown under the experimental conditions described by Wissner *et al.* (2024). Photograph provided by the Laboratory of Microbial Ecology and Marine Natural Products. Chemistry-Sisal Unit, School of Chemistry, Universidad Nacional Autónoma de México (UNAM).

**S2 Fig. BUSCO completeness assessment of the *Streptomyces* sp. NCA360 genome assembly.** BUSCO analysis using the bacterial domain marker dataset (BUSCO- odb_bacteria10) and the *Streptomycetales* group marker dataset (BUSCO- odb_streptomycetales) showed genome completeness of 98.4% and 90.9%, respectively.

**S3 Fig. Genomic coverage and number of BGCs identified in the genome of *Streptomyces* sp. NCA360.** Each point corresponds to a BGC type, positioned according to its total genomic coverage (%). Bubble color and size represent the number of BGCs detected for each type.

**S4 Fig. Structural and regulatory elements present in the nBGCs of the *Streptomyces* sp. NCA360 genome.** The figure shows the structural and regulatory elements identified within the 11 putatively novel biosynthetic gene clusters (nBGCs), including regulatory genes, transcription factor binding sites (TFBSs), and TTA codons.

**S1 Table. Summary of the analysis performed with the Type Strain Genome Server (TYGS).**

**S2 Table. Results of functional genome annotation of *Streptomyces* sp. NCA360 using Prokka v. 1.14.6**

**S3 Table. Information on metabolic pathways related to antibiotic production identified in the genome of *Streptomyces* sp. NCA360**

**S4 Table. Biosynthetic gene clusters identified in the genome of Streptomyces sp. NCA 360, according to antiSMASH v. 7.1.0 results.**

**S5 Table. Resistance model (ResModel) hits identified in the genome of Streptomyces sp. NCA360 (n = 36).**

**S6 Table. Duplicated core genes identified in the *Streptomyces* sp. NCA360 genome (n = 57).**

**S7 Table. Horizontally transferred genes associated with biosynthetic gene clusters in the *Streptomyces* sp. NCA360 genome (16 genes across 18 of 26 BGCs).**

**S8 Table. Genes showing evidence of horizontal gene transfer and proximity to novel biosynthetic gene clusters (nBGCs) in the *Streptomyces* sp. NCA360 genome, identified across 7 of the 11 nBGCs.**

**S9 Table. Information on regulatory genes and transcription factor binding sites identified in novel putative biosynthetic gene clusters associated with antibiotic biosynthesis.**

**S10 Table. FlashPPI results for protein–protein interactions involving 393 proteins encoded within the 11 nBGCs.** Proteins encoded by the 26 identified regulatory genes are highlighted in green.

**S11 Table. FlashPPI results showing 165 protein–protein interactions involving 73 proteins across the 11 nBGCs.**

